# Evidence for the Placenta-Brain Axis: Multi-Omic Kernel Aggregation Predicts Intellectual and Social Impairment in Children Born Extremely Preterm

**DOI:** 10.1101/2020.07.19.211029

**Authors:** Hudson P Santos, Arjun Bhattacharya, Robert M Joseph, Lisa Smeester, Karl CK Kuban, Carmen J Marsit, T. Michael O’Shea, Rebecca C Fry

## Abstract

**Background:** Children born extremely preterm are at heightened risk for intellectual and social impairment, including Autism Spectrum Disorder (ASD). There is increasing evidence for a key role of the placenta in prenatal developmental programming, suggesting that the placenta may explain origins of neurodevelopmental outcomes.

**Methods:** We examined associations between placental genomic and epigenomic profiles and assessed their ability to predict intellectual and social impairment at age 10 years in 379 children from the Extremely Low Gestational Age Newborn (ELGAN) cohort. Assessment of intellectual ability (IQ) and social function was completed with the Differential Ability Scales-II (DAS-II) and Social Responsiveness Scale (SRS), respectively. Examining IQ and SRS allows for studying ASD risk beyond the diagnostic criteria, as IQ and SRS are continuous measures strongly correlated with ASD. Genome-wide mRNA, CpG methylation and miRNA were assayed with the Illumina Hiseq 2500, HTG EdgeSeq miRNA Whole Transcriptome Assay, and Illumina EPIC/850K array, respectively. We conducted genome-wide differential mRNA/miRNA and epigenome-wide placenta analyses. These molecular features were integrated for a predictive analysis of IQ and SRS outcomes using kernel aggregation regression. We lastly examined associations between ASD and the genomically-predicted component of IQ and SRS.

**Results:** Genes with important roles in placenta angiogenesis and neural function were associated with intellectual and social impairment. Kernel aggregations of placental multi-omics strongly predicted intellectual and social function, explaining approximately 8% and 12% of the variance in SRS and IQ scores via cross-validation, respectively. Predicted in-sample SRS and IQ showed significant positive and negative associations with ASD case-control status.

**Limitations:** The ELGAN is a cohort of children born pre-term, andgeneralization may be affected by unmeasured confounders associated with low gestational age. We conducted external validation of predictive models, though the sample size of the out-sample dataset (*N* = 49) and the scope of the available placental datasets are limited. Further validation of the models is merited.

**Conclusions:** Aggregating information from biomarkers within and between molecular data types improves prediction of complex traits like social and intellectual ability in children born extremely preterm, suggesting that traits influenced by the placenta-brain axis may be omnigenic.

## Background

Despite substantial research efforts to elucidate the etiology of neurodevelopmental impairment [1], little is known about genomic and epigenomic factors influencing trajectories of neurodevelopment, such as those associated with preterm delivery [2]. Children born extremely preterm are at increased risk not only for intellectual impairment but also for Autism Spectrum Disorder (ASD) [3,4], often accompanied by intellectual disability. In addition, preterm-born children have consistently been observed to manifest social difficulties (e.g., fewer prosocial behaviors) in childhood and adolecense that do not meet diagnostic criteria for ASD [5].

The placenta is posited as a critical determinant of both immediate and long-lasting neurodevelopmental outcomes in children [1]. The placenta is involved in hormone and neurotransmitter production and transfer of nutrients to the fetus, thus having direct influence on brain development. This connection between the placenta and the brain is termed the placenta-brain axis [6]. Epidemiological and animal studies have linked genomic and epigenomic alterations in the placenta with neurodevelopmental disorders and normal neurobehavioral development [7–9]. For example, the Markers of Autism Risk in Babies: Learning Early Signs (MARBLES) study has identified differentially methylated region containing putative fetal brain enhancer between in placentas from ASD (*N* = 24) and typically developing (n = 23) children [10]. However, identifying genomic signatures of risk for neurodevelopmental disorders such as ASD in placenta is a challenging. Further study of molecular interactions representing the placenta-brain axis may advance our understanding of fetal mechanisms involved in aberrant neurodevelopment [6].

Most prior studies have investigated single molecular levels of the placenta genome or epigenome, precluding analysis of possible interactions that could be linked to neurodevelopmental outcomes. Examining only a single molecular feature, or a single type of features even at a genomic scale can still result in much unexplained variation in phenotype due to potentially important interactions between multiple features [11,12]. This observation is in line with Boyle *et al*.’s omnigenic model [13,14], which proposes that gene regulatory networks are so highly interconnected that a large portion of the heritability of complex traits can be explained by effects on genes outside core pathways. Molecular integration to identify pathways for fetal neurodevelopment in children has been unexplored but may prove to be insightful in associations with complex diseases [15].

We conducted a genome-wide analysis of DNA methylation, miRNA, and mRNA expression in the placenta, examining individual associations with social and intellectual impairment at 10 years of age in children from the Extremely Low Gestational Age Newborn (ELGAN) study [16]. We then combined the genomic and epigenomic data to identify correlative networks of placental genomic and epigenomic biomarkers predictive of social and intellectual impairment as continuous scales, thus allowing us to study neurodevelopmental difficulties beyond the ASD diagnostic categories [17]. To assess the convergent validity of our behavioral findings, we also examined the association of social and intellectual impairment in relation to ASD diagnoses [18]. To our knowledge, this is the first study to use multiple placental molecular signatures to predict intellectual and social impairment, which may inform a framework for predicting risk of adverse neurocognitive and neurobehavioral outcomes in young children.

## Methods

### ELGAN recruitment and study participants

From 2002-2004, women who gave birth at under 28 weeks gestation at one of 14 medical centers across five U.S. states enrolled in the ELGAN study [16]. The Institutional Review Board at each participating institution approved study procedures. Included were 411 of 889 children with both placental molecular analysis and a 10-year follow-up assessment.

### Social and cognitive function and ASD at 10 years of age

Trained child psychologist examiner [5,19] evaluated general cognitive ability (IQ) with the School-Age Differential Ability Scales-II (DAS-II) Verbal and Nonverbal Reasoning subscales [20]. The Social Responsiveness Scale (SRS) was used to assess severity of ASD-related social deficits in 5 subdomains: social awareness, social cognition, social communication, social motivation, and autistic mannerisms [21]. We used the gender-normed T-score (SRS-T; intended to correct gender differences observed in normative samples) as continuous measure of social deficit [22]. All participants were assessed for ASD [18]. Diagnostic assessment of ASD was conducted with three well-validated measures, administered sequentially. First, the Social Communication Questionnaire (SCQ) was administered to screen for potential ASD, using a score ≥ 11 to increase sensitivity relative to the standard criterion score of ≥ 15 [18,23]. For children who screened positive on the SCQ criterion, we conducted the Autism Diagnostic Interview–Revised (ADI-R) with the primary caregiver [24]. All children who met ADI-R criteria for ASD, or who had a prior clinical diagnosis of ASD and/or exhibited symptoms of ASD during cognitive testing according to the site psychologist) were then assessed with the Autism Diagnostic Observation Schedule, Second Version (ADOS-2), which served as the criterion measure of ASD in this study [25]. All ADOS-2 administrations were independently scored by a second rater with autism diagnostic and ADOS-2 expertise. In cases of scoring disagreements, consensus was reached via discussion between raters. Item-by-item inter-rater agreement for the 14 ADOS-2 diagnostic algorithm scores was on average 0.93 (*SD* = 0.12). These developmental assessment procedures and all relevant test scores for ASD and intellectual function are reported in a prior publication [19].

### Placental DNA and RNA extraction

After delivery, placentas were biopsied under sterile conditions. We collected a piece of the chorion, representing the fetal side of the placenta [26]. More specifically, placentas were placed in a sterilized basin and biopsied by pulling back the amnion to expose the chorion at the midpoint of the longest distance between the cord insertion and edge of the placental disk. A sample from the *fetal side* of the placenta was removed by applying traction to the chorion and underlying trophoblast tissue. The specimen was placed in a cryogenic vial and immersed in liquid nitrogen. To preserve DNA and RNA integrity, specimens were stored at −80°C until processed. For processing, a 0.2g subsection of the placental tissue was cut from the frozen biopsy and washed with sterile 1x phosphate-buffered saline to remove any remaining blood. Samples were homogenized using a lysis buffer, and the homogenate was separated into aliquots. This process was detailed in a prior publication [27]. Nucleic acids were extracted from the homogenate using AllPrep DNA/RNA/miRNA Universal kit (Qiagen, Germany). The quantity and quality of DNA and RNA were analyzed using the NanoDrop 1000 spectrophotometer and its integrity verified by the Agilent 2100 BioAnalyzer.

### Epigenome-wide placental DNA methylation

Extracted DNA sequences were bisulfate-converted using the EZ DNA methylation kit (Zymo Research, Irvine, CA) and followed by quantification using the Infinium MethylationEPIC BeadChip (Illumina, San Diego, CA), which measures CpG loci at a single nucleotide resolution, as previously described [26–29]. Quality control and normalization were performed resulting in 856,832 CpG probes from downstream analysis, with methylation represented as the average methylation level at a single CpG site (β-value) [27,30–32]. DNA methylation data was imported into R for pre-processing using the *minfi* package [30]. Quality control was performed at the sample level, excluding samples that failed and technical duplicates; 411 samples were retained for subsequent analyses. Functional normalization was performed with a preliminary step of normal-exponential out-of band (*noob*) correction method [33] for background subtraction and dye normalization, followed by the typical functional normalization method with the top two principal components of the control matrix [31,34]. Quality control was performed on individual probes by computing a detection *P* value and excluded 806 (0.09%) probes with non-significant detection (*P* > 0.01) for 5% or more of the samples. A total of 856,832 CpG sites were included in the final analyses. Lastly, the *ComBat* function was used from the *sva* package to adjust for batch effects from sample plate [83]. The data were visualized using density distributions at all processing steps. Each probe measured the average methylation level at a single CpG site. Methylation levels were calculated and expressed as *β* values (*β* = intensity of the methylated allele (*M*))/(intensity of the unmethylated allele (*U*) + intensity of the methylated allele (*M*) + 100). *β* values were logit transformed to *M* values for statistical analyses [35].

### Genome-wide placental mRNA and miRNA expression

mRNA expression was determined using the Illumina QuantSeq 3’ mRNA-Seq Library Prep Kit, a method with high strand specificity. mRNA-sequencing libraries were pooled and sequenced (single-end 50 bp) on one lane of the Illumina Hiseq 2500. mRNA were quantified through pseudo-alignment with *Salmon* v.14.0 [36] mapped to the GENCODE Release 31 (GRCh37) reference transcriptome. miRNA expression profiles were assessed using the HTG EdgeSeq miRNA Whole Transcriptome Assay (HTG Molecular Diagnostics, Tucson, AZ). miRNA were aligned to probe sequences and quantified using the HTG EdgeSeq System [37]. Genes and miRNAs with less than 5 counts for each sample were filtered, resulting in 11,224 genes and 2,047 miRNAs for downstream analysis. Distributional differences between lanes were first upper-quartile normalized [38]. Unwanted technical and biological variation (e.g. tissue heterogeneity) was then estimated using *RUVSeq* [39], where we empirically defined transcripts not associated with outcomes of interest as negative control housekeeping probes [40]. One dimension of unwanted variation was removed from the variance-stabilized transformation of the gene expression data using the *limma* package [40–43]

### Statistical Analysis

All code and functions used in the statistical analysis can be found at https://github.com/bhattacharya-a-bt/multiomics_ELGAN.

### Correlative analyses between SRS, IQ, and ASD

Associations among SRS scores, IQ and ASD were assessed using Pearson correlations with estimated 95% confidence intervals, and the difference in distributions of SRS and IQ across ASD case-control was assessed using Wilcoxon rank-sum tests. Associations between demographic variables (race, sex, maternal age, number of gestational days, maternal smoking status, placental inflammation, birth weight *Z*-score and mother’s insurance) with SRS and IQ were assessed using multivariable regression, assessing the significance of regression parameters using Wald tests of significance and adjusting for multiple testing with the Benjamini-Hochberg procedure [44].

### Genome-wide molecular associations with SRS and IQ

Once associations between SRS and IQ and ASD were confirmed, we utilized continuous SRS and IQ measures as the main outcomes of interest. Associations between mRNA expression or miRNA expression with SRS and IQ were estimated through a negative binomial linear model using *DESeq2* [43]. Epigenome-wide associations (EWAS) of CpG methylation sites with outcomes were assessed using robust linear regression [45] with test statistic modification through an empirical Bayes procedure [42], described previously [27]. Both the differential mRNA and miRNA expression and EWAS models controlled for the following covariates: race, age, sex, number of gestational age days, birth weight *Z*-score, and education level of the mother. Multiple testing was adjusted for using the Benjamini-Hochberg procedure [44].

### Placental multi-molecular prediction of SRS and IQ

We next assessed how well an aggregate of one or more of the molecular datasets (CpG methylation, mRNA expression, and miRNA expression) predicted continuous SRS and IQ scores. The analytical scheme is summarized in **Figure 1**, using 379 samples with data for all three molecular datasets (DNA methylation, miRNA, and mRNA). Briefly, we first adjusted the outcome variables and molecular datasets for above noted demographic and clinical covariates using *limma* [46] to account for associations between the outcomes and these coviarates in the eventual predictive models. Next, to model the covariance between samples within a single molecular profile, we aggregated the molecular datasets with thousands of biomarkers each into a *molecular kernel* matrix. A *molecular kernel* matrix represents the inter-sample similarities in a given molecular profile (**Supplementary Methods**). A linear or non-linear kernel aggregation may aid in prediction of complex traits by capturing non-additive effects [47–50], which represents a sizable portion of phenotypic variation [51,52]. Using all individual, pairwise, and triplet-wise combinations of molecular kernel matrices, we fitted predictive models of SRS and IQ based on linear mixed modeling [50] or kernel regression least squares (KRLS) [53] and assessed predictive performance with McNemar’s adjusted *R*^2^ via Monte Carlo cross validation [54]. We also optimized predictive models for the number of included biomarkers per molecular profile. Extensive model details, as well as alternative models considered, are detailed in **Supplemental Methods.**

**Figure 1:**
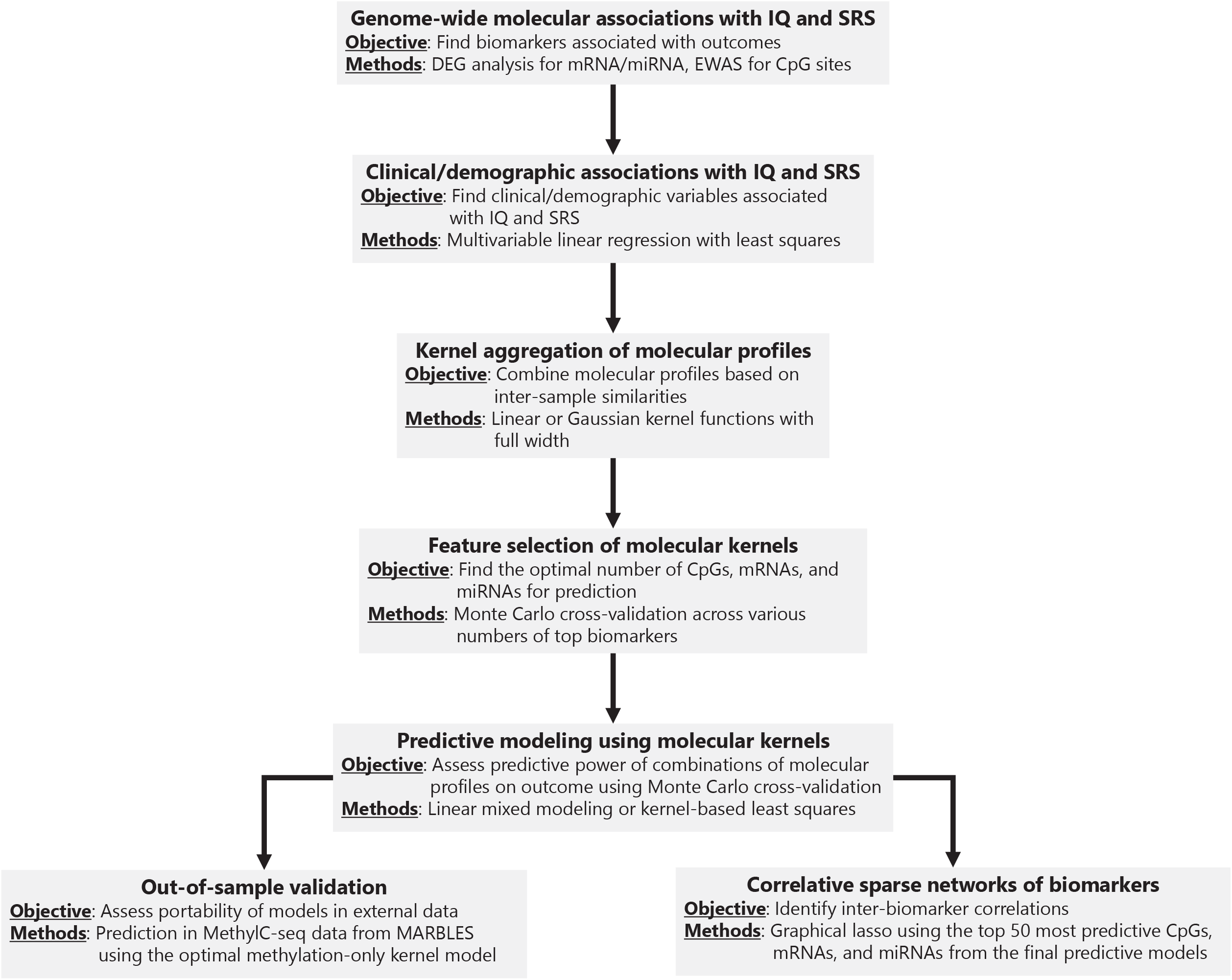
Scheme for kernel aggregation and prediction models. (1) Design matrices for CpG sites, mRNAs, and miRNAs are aggregated to form a linear or Gaussian kernel matrix that measures the similarity of samples. (2) Clinical variables are regressed out of the outcomes IQ and SRS and from the omic kernels to limit influence from these variables. (3) Using 50-fold Monte Carlo cross-validation on 75%-25% training-test splits, we train prediction models with the kernel matrices for IQ and SRS in the training set and predict in the test sets. Prediction is assessed in every fold with adjusted *R*^2^ and averaged for an overall prediction metric.

### Validation in external dataset

Lack of studies that consider placental mRNA, CpG methylation and miRNA data with long-term child neurodevelopment limit the ability to extablish external validation. We obtained one external placental CpG methylation dataset from the Markers of Autism Risk in Babies-Learning Early Signs (MARBLES) cohort [10]. To assess out-of-sample performance of kernel models for methylation, we downloaded MethylC-seq data for 47 placenta samples, 24 of which identified as ASD cases (NCBI Gene Expression Omnibus accession numbers GSE67615) [10]. *β*-values for DNA methylation were extracted from BED files and transformed into *M*-values with an offset of 1 [35], and used the best methylation-only predictive model to predict SRS and IQ in these 47 samples, as detailed in **Supplemental Methods**.

### Correlative networks

In the final KRLS predictive models for both IQ and SRS including all three molecular profiles, we extracted the top 50 most predictive (largest point-wise effect sizes) CpGs, miRNAs, and mRNAs of SRS and IQ. A sparse correlative network was inferred among these biomarkers that links biomarkers based on the strength of correlative signals using graphical lasso in *qgraph* [55,56].

## Results

### SRS and IQ are well associated with ASD

Although the sample is enriched for ASD cases (*N* = 35 cases, 9.3% of the sample) relative to non-preterm cohorts, there is still a relatively low case-control ratio for a genome-wide study of this sample size (descriptive statistics for relevant covariates in **Table 1**). Therefore, we considered continuous measures of social impairment (SRS) and cognitive development (IQ) at age 10 for both associative and predictive analyses. Using continuous variables for SRS and IQ allow us to to study complexities beyond the ASD diagnostic categories [15,17]. **Figure 2A-B** shows the relationship between SRS, IQ, and ASD. The mean SRS is significantly higher in ASD cases compared to controls (mean difference of 1.74, 95% *CI*: (1.41,2.07)). Mean IQ is significantly lower in ASD cases versus controls (mean difference of – 2.23, 95% *CI* (−2.46,-1.96)). Furthermore, SRS and IQ are negatively correlated (Pearson *ρ* = −0.47,95% *CI*: (−0.55, −0.39)). We also measured associations between demographic characteristics with SRS and IQ (**Figure 2C**) using multivariable regression. Male sex is associated with lower IQ, while public health insurance is associated with both lower IQ and increased social impairment. Demographic variables included in the multivariable regression explain approximately 12% and 15% of the total variance explained in IQ and SRS, as measured by adjusted *R*^2^, with a summary of regression parameters in **Table 2**. Based on the associations identified here and the value of inclusion of continuous measures, subsequent genomic and epigenomic analyses control for demographic covariates.

**Figure 2:**
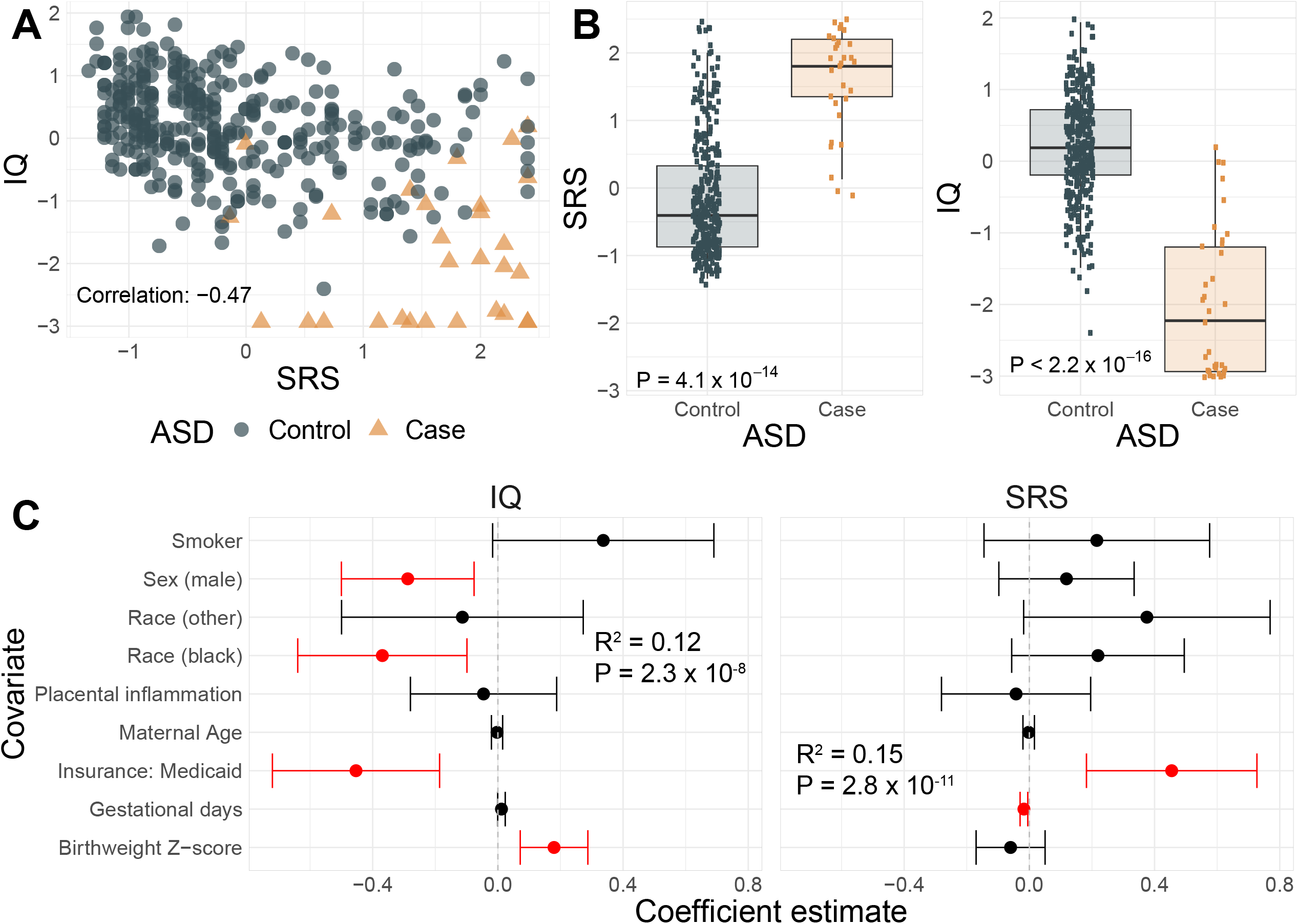
Associations between SRS, IQ, and ASD and with clinical variables. **(A)** Scatter plot of SRS (X-axis) and IQ (Y-axis) colored by ASD case (orange) and control (blue) status. **(B)** Boxplots of SRS and IQ across ASD case-control status. *P*-value from a two-sample Mann-Whitney test is provided. **(C)** Caterpillar plot of multivariable linear regression parameters of IQ and SRS using clinical variables. Points give the regression parameter estimates with error bars showing the 95% FDR-adjusted confidence intervals [44]. The null value of 0 is provided for reference with the dotted line.

**Table 1:**
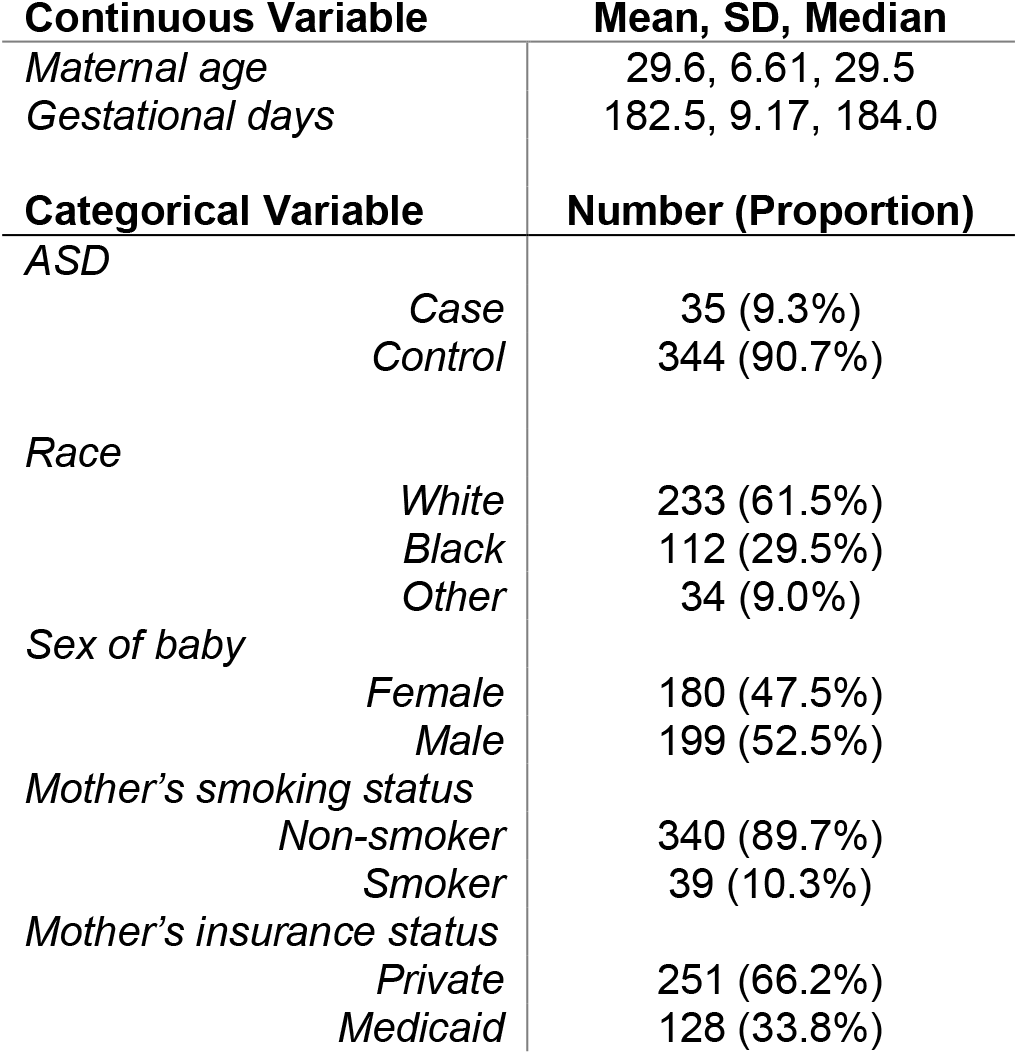
Descriptive statistics for demographi and clinical covariates.

| Continuous Variable |  | Mean, SD, Median |
| --- | --- | --- |
| Maternal age |  | 29.6, 6.61, 29.5 |
| Gestational days |  | 182.5, 9.17, 184.0 |
| Categorical Variable |  | Number (Proportion) |
| ASD |  |  |
|  | Case | 35 (9.3%) |
|  | Control | 344 (90.7%) |
| Race |  |  |
|  | White | 233 (61.5%) |
|  | Black | 112 (29.5%) |
|  | Other | 34 (9.0%) |
| Sex of baby |  |  |
|  | Female | 180 (47.5%) |
|  | Male | 199 (52.5%) |
| Mother's smoking status |  |  |
|  | Non-smoker | 340 (89.7%) |
|  | Smoker | 39 (10.3%) |
| Mother's insurance status |  |  |
|  | Private | 251 (66.2%) |
|  | Medicaid | 128 (33.8%) |

**Table 2:**
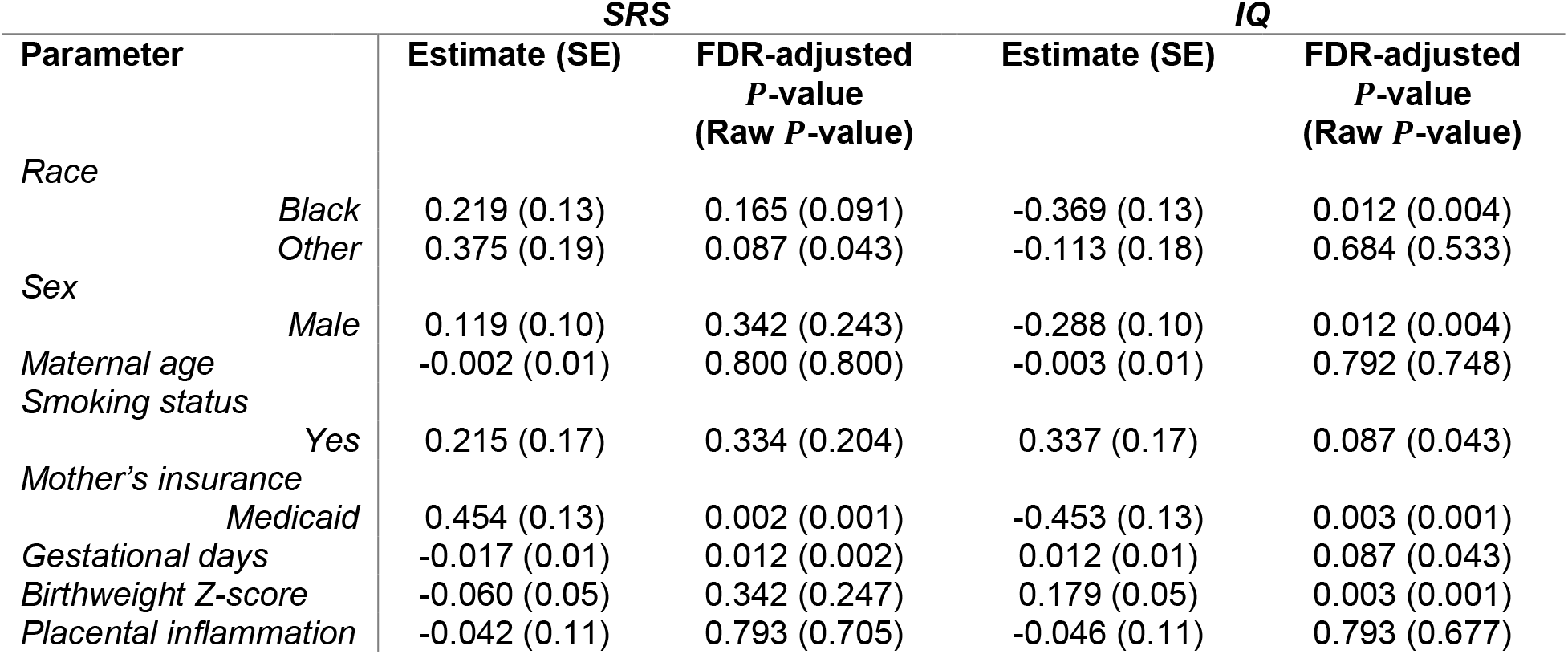
Summary of regressions of SRS and IQ against clinical covariates.

### Genome-wide associations of mRNA, miRNA, and CpGs with SRS and IQ

Genome-wide association tests between each of the individual placental molecular datasets (e.g. the placental mRNA data, the CpG methylation, or the miRNA datasets) in relation to SRS and IQ (see **Methods**) identified two genes with mRNA expression significantly associated with SRS at FDR-adjusted *P* < 0.01 (Hdc Homolog, Cell Cycle Regulator [*HECA*], LIM Domain Only 4 [*LMO4*]). We did not find CpG sites or miRNAs associated with SRS (**Table 3**). Associations between IQ and the mRNA expression, at FDR-adjusted *P* < 0.01, were observed at four genes, namely Ras-Related Protein Rab-5A (*RAB5A*), Transmembrane Protein 167A (*TMEM167A*), Signal Transducer and Activator of Transcription 2 (*STAT2*), ITPRIP Like 2 (*ITPRIPL2*). One CpG site (*cg09418354* located in the gene Carbohydrate Sulfotransferase 11 (*CHST11*) displayed an association with IQ, and no miRNAs were associated with IQ (**Table 3**). Manhattan plots (**Supplemental Figure 1**) show the strength of associations of all biomarkers by genomic position. Summary statistics for these associations are provided in **Supplemental Materials**. No mRNAs, CpG sites, or miRNAs were significantly associated with both SRS and IQ, though effect sizes for associations with the same features were in opposite directions (see **Supplemental Materials**).

**Table 3:** Summary of genome-wide associations of molecular profiles with SRS and IQ at FDR-adjusted *P* < 0.01.

| **SRS** |  |  |
| --- | --- | --- |
| **Biomarker** | **Effect size** | **FDR-adjusted P-value** |
| *mRNA expression* |  |  |
| *HECA* | 0.571 | 0.001 |
| *LMO4* | 0.467 | 0.001 |
| **IQ** |  |  |
| **Biomarker** |  |  |
| *mRNA expression* |  |  |
| *RAB5A* | -0.516 | 0.002 |
| *TMEM167A* | -0.632 | 0.004 |
| *ITPRIPL2* | -0.557 | 0.004 |
| *STAT2* | -0.584 | 0.004 |
| *CpG methylation site* |  |  |
| *cg09418354* | -0.005 | 0.002 |

### Kernel regression shows predictive utility in aggregating multiple molecular datasets

Because the genome wide association analyses revealed few mRNAs, CpG sites or miRNAs that were associated with SRS or IQ with large effect sizes, we next assessed the impact of aggregating these molecular datasets on prediction of SRS and IQ. This was done to account for the considerable number of biomarkers that have moderate effect sizes on outcome. To find the most parsimonious model with the greatest predictive performance, we first selected the optimal number of biomarkers per molecular profile for each outcome that gave the largest mean adjusted *R*^2^ in predictive models with only one of the three molecular datasets (see **Supplemental Methods**). **Figure 3A** shows the relationship between the number of biomarkers from the mRNA expression, CpG level, miRNA expression datasets and their predictive performance. In general, predictive performance steadily increased as the number of biomarker features increased until reaching a tipping point where predictive performance decreased (**Figure 3A**). Overall, for CpG methylation, the top (lowest P-values of association) 5,000 CpG features showed the greatest predictive performance, and for the mRNA and miRNA expression datasets, the top 1,000 features showed the greatest predictive performance.

**Figure 3:**
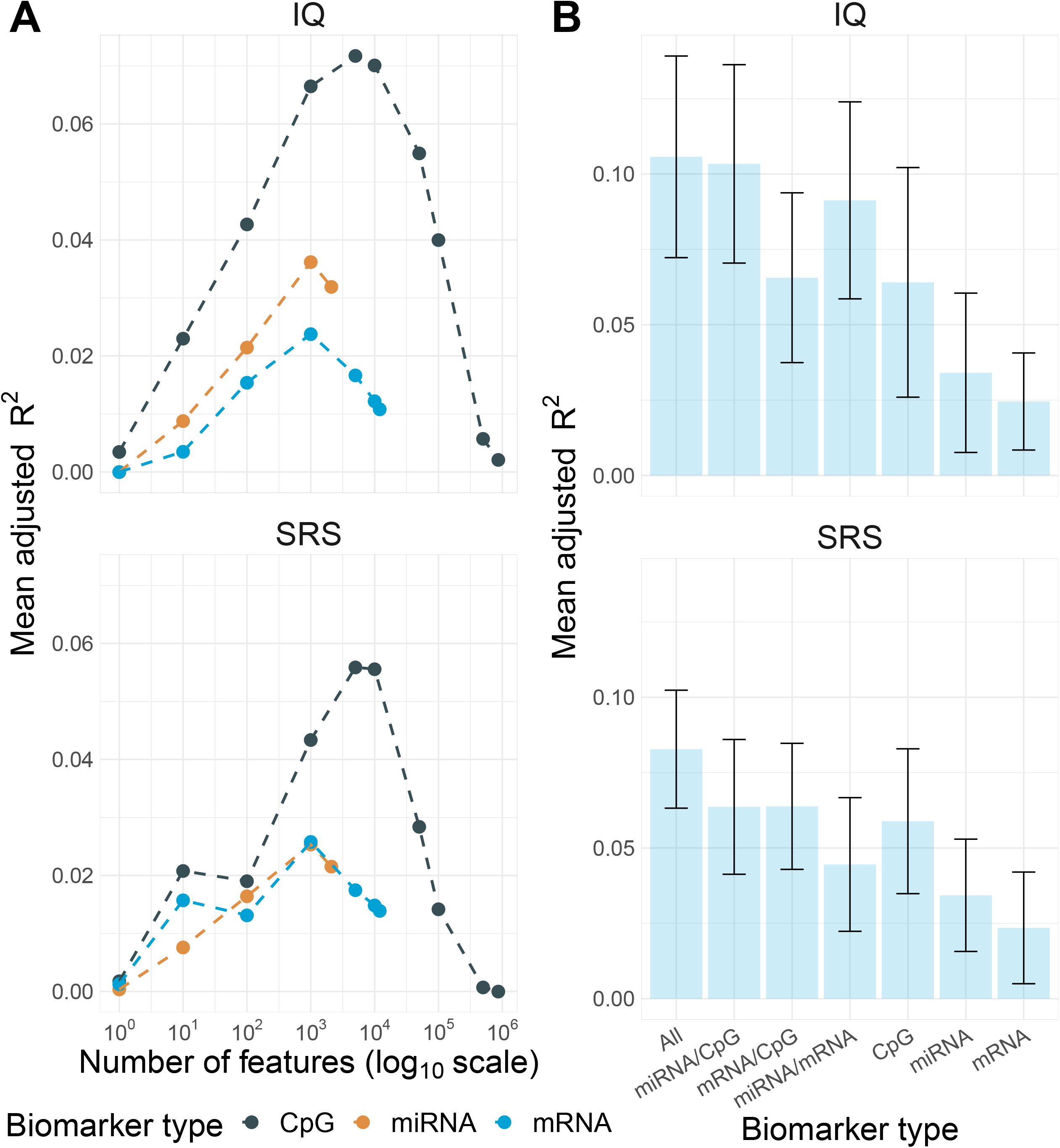
In-sample predictive performance of kernel models. (A) Adjusted mean *R*^2^ (Y-axis) of best kernel models over various numbers of the top biomarkers (X-axis) in the CpG (dark blue), miRNA (orange), and mRNA (light blue) omics over 50 Monte Carlo folds. The X-axis scale is logarithmic. (B) Bar plots of adjusted mean *R*^2^ (Y-axis) for optimally tuned kernel predictive models using all combinations of omics (X-axis) over 50 Monte Carlo folds. The error bar gives a spread of one standard deviation around the mean adjusted *R*^2^.

Using the fully-tuned 7,000 biomarkers (5,000 for CpG methylation and 1,000 for both mRNA and miRNA expression) per molecular dataset with feature selection done in the training set, we trained predictive models (both linear and Gaussian kernel models) using all individual, pair-wise, and triplet-wise combinations of the three molecular datasets. **Figure 3B** shows that whereas the mRNA had the lowest predicted performance to both IQ (*R*^2^ = 0.025) and SRS (*R*^2^ = 0.025), aggregating the mRNA expression, CpG methylation and miRNA expression datasets tends to increase the predictive performance. Specifically, in relation to both outcomes (SRS and IQ), the model using all three integrated datasets shows the greatest predictive performance (mean adjusted *R*^2^ = 0.11 in IQ and *R*^2^ = 0.08 in SRS).

### Correlative networks of placental biomarkers

To gain further understanding of the associations among the identified mRNA, CpG and miRNA biomarkers in the context of IQ and SRS, we extracted (*n* = 50) mRNA, CpGs, and miRNAs that have the largest effect sizes on IQ and SRS in the kernel regression models and inferred sparse correlative networks using the graphical lasso [55,56] (see **Methods**). In the networks (**Supplemental Figure 2**), each molecular dataset clusters by itself, with minimal nodes extending between molecular datasets, and more interconnection is observed between miRNAs and CpG methylation versus mRNAs. These networks point to genes that play important roles in placental angiogenesis and neural function, such as *SMARCA2* (SWI/SNF Related, Matrix Associated, Actin Dependent Regulator Of Chromatin, Subfamily A, Member 2), *SLIT3* (Slit Guidance Ligand 3), and *LZTS2* (Leucine Zipper Tumor Suppressor 2) that have been previously associated with neurodevelopmental disorders, including intellectual disability, social impairment, mood disorders, and ASD [57–62].

### Validation of in-sample and out-sample SRS and IQ prediction with ASD case and control

To contextualize our predictions, we tested whether the predicted SRS and IQ scores generated by our kernel models are associated with ASD case-control status; these predicted SRS and IQ scores represent the portion of the observed SRS and IQ values that our models can predict from placental genomic features. We used the optimal 7,000 biomarker features identified with a 10-fold cross-validation process, splitting samples into 10 hold-out sets and using the remaining samples as a training set to predict SRS and IQ for all 379 samples. After accounting for covariates, the predicted SRS and IQ values from the biomarker data were well-correlated with the observed clinical SRS and IQ values, explaining approximately 8% (approximate Spearman *ρ* = 0.29, cross-validatation *R*^2^ P-value *P* = 7.5 × 10^−9^) and 12% (Spearman *ρ* = 0.35, *P* = 3.6 × 10^−12^) of the variance in the observed SRS and IQ variables, respectively. In addition, we found strong association between the predicted SRS and IQ with ASD case and controls, mean difference of −0.56 (test statistc *W* = 8121,*P* = 6.6 × 10^−4^) for IQ, and mean difference of 0.33 (*W* = 4717, *P* = 0.03) for SRS (**Figure 4**).

**Figure 4:**
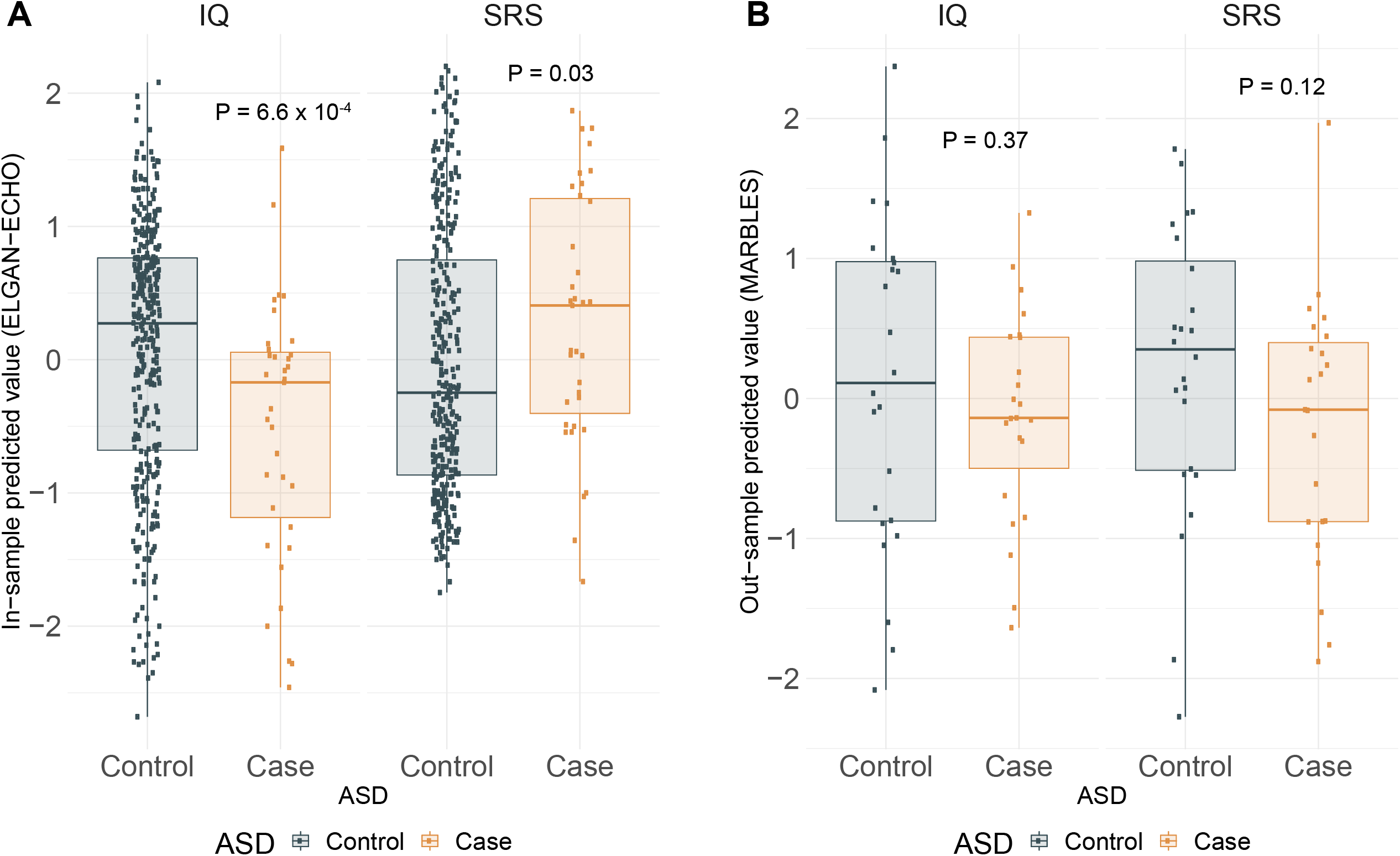
Association of ASD case/control status with predicted SRS and IQ. **(A)** Box-plots of insample predicted IQ (left) and SRS (right) over ASD case/control in ELGAN over 10-fold cross-validation. **(B)** Box-plots of out-sample predicted IQ (left) and SRS (right) over ASD case/control in MARBLES external validation dataset. P-values presented as from a Mann-Whitney test of differences across the ASD case/control groups.

Because we lacked an external dataset with all three molecular data (mRNA, CpG methylation, and miRNA) and cognitive, social impairment and ASD data, we assessed the out-of-sample predictive performance of the CpG methylation-only models using MethylC-seq data from the MARBLES cohort (GEO GSE67615) [10]. We computed predicted IQ and SRS values for 47 placental samples (24 cases of ASD) and assessed differences in mean predicted IQ and SRS across ASD case and control groups. The direction of the association is similar to our data for IQ yet the differences in mean predicted IQ (−0.22, *P* = 0.37) and SRS (−0.42, *P* = 0.12) across ASD groups in MARBLES is not significant (**Figure 4**). This external validation provides some evidence of the portability of our models and merits further future validation of these models, as more placental multi-omic datasets are collected.

## Discussion

We evaluated the predictive capability of three types of genomic and epigenomic molecular biomarkers (mRNA, CpG methylation, and miRNA) in the placenta on cognitive and social impairment in relation to ASD at 10 years of age. Genes that play important roles in placenta angiogenesis and neural function were associated with SRS and IQ. The multi-omic predictions of SRS and IQ are strong and explain up to 8% and 12% of the variance in the observed SRS and IQ variables in 5-fold cross-validation, respectively. This study supports the utility of aggregating information from biomarkers within and between molecular datasets to improve prediction of complex neurodevelopmental outcomes like social and intellectual ability, suggesting that traits on the placenta-brain axis may be omnigenic.

Several genes with known ties to neurodevelopmental disorders distinguished individuals with and without intellectual and social impairmenats. For example, CpG methylation in *SLIT3* was associated with intellectual (IQ) disability. *SLIT3* is highly expressed in trophoblastic endothelial cells [63] and plays a critical role in placental angiogenesis and in the development of neuronal connectivity. Human and animal genetic studies support that *SLIT3* is associated with mood disorders, IQ, and ASD [61,64–66]. *LZTS2*, another gene we found to be associated with IQ, is involved in regulating embryonic development by the *Wnt* signaling pathway [67,68]. Genetic and miRNA expression studies have linked *LZTS2* to social impairment and ASD [69–71]. Furthermore, *LZTS2* is bound by the Chromodomain Helicase DNA Binding Protein 8 gene (*CHD8*), which is associated with brain development in mice and neurodevelopmental disorders in humans [72–74]. In relation to social impairment, *ADAMTS6* was found to be associated with SRS.The *ADAMTS6* gene is a member of the ADAMTS protein family and is regulated by the cytokine TNF-alpha [75]. In previous studies, *ADAMTS6* has been implicated in intellectual disability and growth development and with socially affected traits in pigs [76,77].

Looking into the individual molecular datasets, DNA methylation effects showed the strongest prediction of both SRS and IQ impairment. There is strong evidence suggesting inverse correlation between DNA methylation of the first intron and gene expression across tissues and species [78]. We found that many of the CpG loci with the largest effect sizes on SRS and IQ identified in our analysis are located near DNAase hyperactivity or active regulatory elements for the placenta [79,80], suggesting that these loci likely play regulatory functions. Experimental studies have demonstrated regions of the genome in which DNA methylation is causally important for gene regulation and those in which it is effectively silent [81]. We found that aggregating biomarkers within and between molecular datasets improves prediction of social and cognitive impairment. Specifially, this observation suggests new possibilities to the discovery of candidate genes in the placenta that convey neurodevelopmental risk, improving the understanding of the placenta-brain axis. Recent work in transcriptome-wide association studies (TWAS) are a promising tool that aggregates genetics and transcriptomics to identify candidate trait-associated genes [82,83]. Incorporating information from regulatory biomarkers, like transcription factors and miRNAs, into TWAS increases study power to generate hypotheses about regulation [84,85]. Given our observations in this analysis and the number of the integrated molecular datasets, we believe that the ELGAN study can be used to train predictive models for placental transcriptomics from genetics, enriched for regulatory elements [85]. These transcriptomic models can then be applied to genome-wide association study cohorts to study the regulation of gene-trait associations in the placenta.

### Limitations

When interpreting the results of this study, some factors should be considered. Extremely preterm birth is strongly associated with increased risk for neurodevelopmental disorders [18]. This association may lead to bias in estimated associations between the molecular biomarkers and outcomes, especially when unmeasured confounders are linked to both pre-term birth and autism [86]. Still, to our knowledge the ELGAN cohort is currently the largest available placental repository with both multiple molecular datasets and long-term neurodevelopmental assessment of the children. Second, as the placenta is comprised of several heterogeneous cell types, tissue-specific molecular patterns in the placenta should be taken into consideration when interpreting these findings in relation to other tissue samples; future comparison between tissues will not be straightforward. Lastly, to test the reproducibility and robustness of our kernel models, we believe further out-of-sample validation is required, using datasets with larger sample sizes and similar molecular datasets. Though in-sample predictive performance is strong, platform differences between the ELGAN training set (assayed with EPIC BeadChip) and validation set (MethylC-seq) may lead to loss of predictive power. As our optimal models all aggregate various datasets, the dearth of data for the placenta, in the context of social and intellectual impairment, makes out-of-sample validation especially challenging. Lack of external validation may render our analysis exploratory in nature, but we provide evidence of a link between molecular features within the fetal placenta and social and cognitive outcomes in children that merits future investigation.

### Conclusions

Our analysis underscores the importance of synthesizing data representing various levels of biological data to understand distinct genomic and epigenomc underpinnings of complex developmental deficits, like intellectual and social impairment. This study provides novel evidence for the omnigenicity of the placenta-brain axis in the context of social and intellectual impairment.

## Abbreviations

(ELGAN): Extremely Low Gestational Age Newborn
(IQ): Intellectual ability
(DAS-II): Differential Ability Scales-II
(SRS): Social Responsiveness Scale
(ASD): Autism Spectrum Disorder
(SRS-T): SRS gender-normed T-score
(EWAS): Epigenome-wide association study
(MARBLES): Markers of Autism Risk in Babies-Learning Early Signs
(KRLS): Kernel regression least squares

## Declarations

### Ethics approval and consent to participate

The study was approved by the Institutional Review Board of the University of North Carolina at Chapel Hill. All participants consented to the study as per IRB protocol.

### Consent for publication

Not applicable

### Availability of data and materials

Multiomic data from the ELGAN study is available from the NCBI Gene Expression Omnibus GSE154829. All genomic and clinical data is also available upon request to H.P.S. For validation, we used MethylC-seq data from the MARBLES study available at GSE67615.

## Authors’ Contributions

H.S.P, R.M.J., L.S., K.C.K.K., C.J.M., T.M.O, and R.C.F. conceived and designed the study. H.S.P. and A.B. acquired and analyzed the data. H.S.P., A.B. and R.C.F. interpreted data. H.S.P. and A.B. drafted the work and all authors revised. All authors have approved the submitted version and are accountable for their own contributions.

## Acknowledgements

We would like to thank the study participants of the ELGAN-ECHO study. We would also like to thank Michael Love and Lana Garmire for helpful discussion during the research process.

