## Supplemental Materials for "Evidence for the Placenta-Brain Axis: Multi-Omic Kernel Aggregation Predicts Intellectual and Social Impairment in Children Born Extremely Preterm"

Supplementary Methods

*Projection to kernel space*

For a given omic design matrix $\boldsymbol{X}_m$ on $n$ samples and $p_m$ features, with each column standard-normalized to zero mean and unit variance, we project the omic into a sparse kernel space using the simple linear kernel:

$$\boldsymbol{K}_{\boldsymbol{linear}_m} = p_m^{-1}\boldsymbol{X}_m\boldsymbol{X}_m^T,$$

resulting in $\boldsymbol{K}_{\boldsymbol{linear}_m}$, an $n \times n$ matrix. We call $\boldsymbol{K}_{\boldsymbol{linear}_m}$ the omic linear kernel matrix for omic $m$.

Alternatively, for a given omic $\boldsymbol{X}_m$ on $n$ samples and $p_m$ features, with each column standard-normalized to zero mean and unit variance, we project the omic design matrix into a sparse Gaussian kernel space using the Gaussian kernel function. For samples $i$ and $j$, corresponding to the $i$th and $j$th rows of $\boldsymbol{X}_m$, the $i,j$-th element of the Gaussian kernel matrix $\boldsymbol{K}_{\boldsymbol{Gauss}_m}$ for omic $m$ is defined as

$$\boldsymbol{K}_{\boldsymbol{Gauss}_m} = \exp\left(\frac{-\|x_{i.}-x_{j.}\|^2}{p_m}\right).$$

To compare two omic kernels $\boldsymbol{K}_{m_1}$and $\boldsymbol{K}_{m_2}$ in the same kernel space (i.e. two linear kernel matrices or two Gaussian kernel matrices), we compute their kernel alignment $S$, a measure of similarity between two omic kernels [7] using the standardized Frobenius inner product of two matrices:

$$S = \frac{\langle \boldsymbol{K}_{m_1}, \boldsymbol{K}_{m_2}\rangle_F}{\sqrt{\langle \boldsymbol{K}_{m_1}, \boldsymbol{K}_{m_1}\rangle_F \langle \boldsymbol{K}_{m_2}, \boldsymbol{K}_{m_2}\rangle_F}},$$

where, for two $n \times n$ matrices $\boldsymbol{A}$ and $\boldsymbol{B}$, $\langle \boldsymbol{A}, \boldsymbol{B}\rangle_F = \sum_{i,j=1}^n A_{ij}B_{ij}$.

*Multiomic kernel predictive model with linear kernels*

We consider the following linear model, assuming $M$ total omics considered:

$$\boldsymbol{Y} = \boldsymbol{X}_c\boldsymbol{\beta}_C + \boldsymbol{U} + \boldsymbol{\epsilon},$$

where $\boldsymbol{Y}$ is an outcome of interest (either SRS or IQ), standardized to zero mean and unit variance, $\boldsymbol{X}_C$ is the design matrix of covariates, $\boldsymbol{\beta}_C$ is a vector of fixed effects for covariates, $\boldsymbol{U}$ is a vector of omic predictive scores, and $\epsilon$ is Gaussian random error with zero mean and identity covariance matrix. We assume that the omic predictive scores are normally distributed with zero mean and a fused kernel matrix as its variance-covariance matrix, i.e. $\boldsymbol{U} \sim N(0, \boldsymbol{K})$ and $\boldsymbol{K} = \sum_{m=1}^{M} \boldsymbol{K}_m$. This model is simply a traditional linear mixed model, treating clinical covariates as fixed effects and aggregating all omics into omic predictive scores that are treated as random effects clustered around zero. It is straightforward that

$$\boldsymbol{U} = \sum_{m=1}^{M} \boldsymbol{U}_m = \sum_{m=1}^{M} \sum_{j=1}^{p_m} \boldsymbol{X}_{mj} \eta_{mj},$$

where $\boldsymbol{U}_m$ is the vector of omic predictive scores for the $m$th omic profile and the weight sum of omic features $\boldsymbol{X}_{mj}$ with random effects coefficients $\eta_{mj}$. Assuming $\eta_{mj} \sim N(0, {\sigma_m^2}/{p_m})$, we see that $\boldsymbol{U}_m \sim N(0, \sigma_m^2 \boldsymbol{K}_m)$ and $\boldsymbol{U} \sim N(0, \boldsymbol{K})$.

*Kernel modeling training and evaluation for linear kernels*

We evaluate our models through 50-fold Monte Carlo cross-validation, using methods similar to Zhu et al [8]. Given a single cross-validation fold, with 75% of the data in training set and 25% in a test set, we can divide $\boldsymbol{Y} = \begin{bmatrix} \boldsymbol{Y}_{train} \\ \boldsymbol{Y}_{test} \end{bmatrix}$ and $\boldsymbol{K}_{\boldsymbol{linear}}$ into four blocks:

$$\boldsymbol{K}_{\boldsymbol{linear}} = \begin{bmatrix} \boldsymbol{K}_{train} & \boldsymbol{K}_{cov} \\ \boldsymbol{K}_{cov}^T & \boldsymbol{K}_{test} \end{bmatrix},$$

where $\boldsymbol{K}_{train}$ is the variance matrix of the training set, $\boldsymbol{K}_{test}$ is the variance matrix of the test set, and $\boldsymbol{K}_{cov}$ is the covariance matrix between the training and test sets. We find the maximum likelihood estimator $\widehat{\boldsymbol{\beta}}_C$ for the fixed effects, the best linear unbiased predictor $\widehat{\boldsymbol{U}}_{train}$ of the omic predictive scored of the training set, and the restricted maximum likelihood estimators $\hat{\sigma}_m^2$ for the variance components of the fused kernel, using *rrBLUP* [9]. We then estimate the total predictive scores $\widehat{\boldsymbol{Y}}_{test}$:

$$\widehat{\boldsymbol{Y}}_{test} = \boldsymbol{X}_{C_{test}} \widehat{\boldsymbol{\beta}}_C + \boldsymbol{K}_{cov} \boldsymbol{K}_{train}^{-1} \widehat{\boldsymbol{U}}_{train}$$

We then compute the adjusted $R^2$ between the $\boldsymbol{Y}_{test}$ and $\widehat{\boldsymbol{Y}}_{test}$ to assess the predictive performance of the omic predictive scores. This process is repeated across 100 folds and the adjusted $R^2$ are averaged to create a predictive index for a given clinical and omic profile. We consider all possible combinations of omics after regressing out clinical covariates in these multiomic kernel models. External validation is conducted similarly; however, as the relevant clinical covariates were not available in the MARBLES dataset, we do not adjust the molecular profiles for these clinical covariates in the predictive models applied to this dataset[10].

*Kernel regularized least squares regression for Gaussian kernels and model evaluation*

We employ a similar Monte Carlo cross-validation scheme, splitting our data into 75%-25% training and test sets across 50 folds. We implement kernel-based regularized squares regression through the *KRLS* package [11], that minimizes the Tikhonov objective function over squared loss. Briefly, the objective of KRLS, in our case, is to find the $c$ that minimizes

$$T(c) = \sum_{i=1}^{n}\left(\boldsymbol{Y} - \boldsymbol{K}_{\boldsymbol{Gauss}_m}c\right)^T(Y - \boldsymbol{K}_{\boldsymbol{Gauss}_m}c) + \lambda c^T\boldsymbol{K}_{\boldsymbol{Gauss}_m}c,$$

where $\lambda$ is tuned via leave-one-out cross-validation. Furthermore, KRLS can compute the pointwise partial derivatives of the fitted function with respect to each predictor using estimators developed by Hainmuller and Hazlett [11]. These pointwise partial derivatives can be used to examine the marginal effect of every feature in the omic design matrix on the outcome of interest, similar to an ordinary least squares regression parameter estimate.

Using the best parameter estimates from the training set, a Gaussian kernel matrix can be computed from the test set, and we define predicted values in the test set as

$$\widehat{\boldsymbol{Y}}_{test} = \boldsymbol{K}_{\boldsymbol{Gauss}_{m_{test}}}\hat{c}.$$

We compute adjusted $R^2$ between the observed and predicted values of the outcome of interest here, as well, to assess predictive performance.

*Sparse regression model and evaluation*

We also consider linear models to predict outcomes of interest using a regularized regression model:

$$\boldsymbol{Y} = \boldsymbol{X_m}\boldsymbol{\beta}_m + \boldsymbol{\epsilon},$$

where $\boldsymbol{X}_m$ and $\boldsymbol{\beta}_m$ are the $n \times p_m$ design matrix and effects for a given omic profile $m$, using Monte Carlo cross-validation to evaluate predictive performance across 100 80%-20% training-test set folds. In the training set, using *glmnet*, we estimate $\widehat{\boldsymbol{\beta}}_m$ using elastic net regularized regression with a mixing parameter of 0.5 (even mixture of LASSO and ridge penalties) and the regularization penalty parameter tuned over 5 folds [12]. Using these parameter estimates, we predict on the test set and evaluate predictive performance using adjusted $R^2$.

*Feature selection*

To tune the number of features to include for each omic in the final predictive models, we tuned over various $P$-value thresholds from fold-wise one-way tests of associations. We split the data into 5 80%-20% training set-test set folds, conducted one way tests of associations in the training set (i.e. differential expression analysis for mRNA and miRNA expression, EWAS for DNA methylation) , and selected all biomarkers with associations with the outcome of interest with $P$-value under a given threshold. Using these biomarkers, we predict the outcome of interest in the test set using both kernel regression methods and compute the adjusted $R^2$ to assess predictive performance.

*External validation using MARBLES dataset*

We obtained one external placental CpG methylation dataset from the Markers of Autism Risk in Babies-Learning Early Signs (MARBLES) cohort [10]. To assess out-of-sample performance of kernel models for methylation, we downloaded MethylC-seq data for 47 placenta samples, 24 of which identified as ASD cases (NCBI Gene Expression Omnibus accession numbers GSE67615) [10]. We extracted $\beta$-values for DNA methylation from BED files and transformed into $M$-values with an offset of 1 [13]. We then used the best linear kernel and kernel regression models to predict SRS and IQ in the MARBLES dataset, as detailed above. It is important to note that not all CpG sites used in the best-methylation model from ELGAN were assayed in the MARBLES external validation set (only approximately 85% overlap). Furthermore, the MARBLES dataset does not have measures of SRS or IQ. Thus, to assess the validity of predicted SRS and IQ estimates in MARBLES, we tested for association between the predicted SRS and IQ values and ASD case-control status.

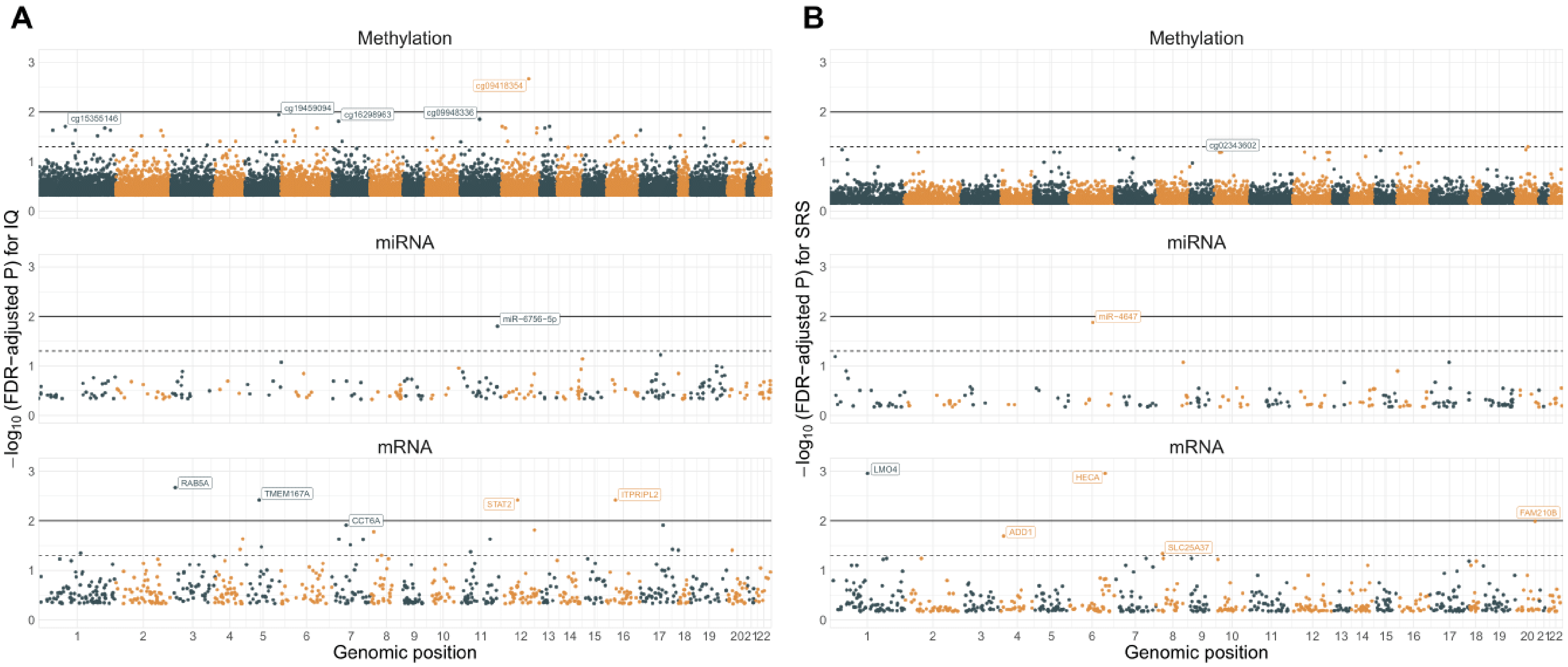


*Supplemental Figure 1:* ***Manhattan plots of one-way omic association tests for IQ and SRS.*** Manhattan plots for one-way tests of association for methylation (top), miRNA (middle), and mRNA expression with IQ (A) and SRS (B). The $\boldsymbol{X}$-axis plots the genomic position of the biomarker and the $\boldsymbol{Y}$-axis plots the Benjamini-Hochberg FDR-adjusted $P$-value for the association with the given outcome. The dotted line provides a reference of FDR-adjusted $P = 0.05$, and the solid line provides a reference of FDR-adjusted $P = 0.01$. Biomarkers are labelled with their association has FDR-adjusted $P = 0.01$.

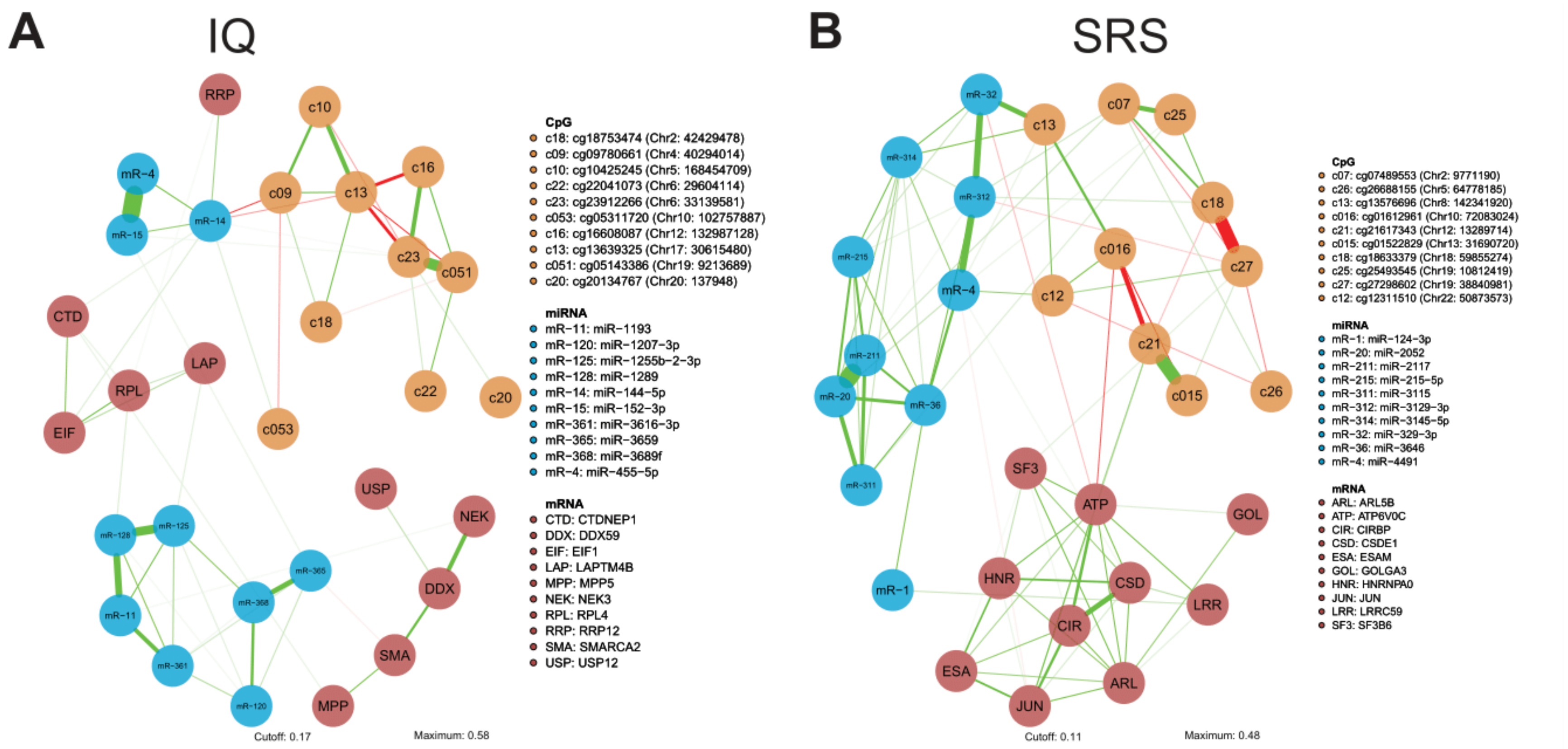


*Figure 2:* ***Sparse correlative networks between top predictive biomarkers of IQ and SRS***. Using the top 50 CpGs (orange nodes), top 50 miRNAs (blue), and top 50 mRNAs (red) that are predictive of IQ (left) and SRS (right), we inferred sparse correlative networks [14, 15]. Nodes are biomarkers and edges show correlations between biomarkers. Positive correlations are shown in green, negative correlations are shown in red, and the thickness of the edge gives the absolute magnitude of correlation.

11. Hainmueller J, Hazlett C. Kernel Regularized Least Squares: Reducing Misspecification Bias with

a Flexible and Interpretable Machine Learning Approach. Polit Anal. 2014;22:143–168.

12. Friedman J, Hastie T, Tibshirani R. Regularization Paths for Generalized Linear Models via Coordinate Descent. J Stat Softw. 2010;33:1–22.

13. Du P, Zhang X, Huang C-C, Jafari N, Kibbe WA, Hou L, et al. Comparison of Beta-value and M-value methods for quantifying methylation levels by microarray analysis. BMC Bioinformatics. 2010;11:587.

14. Epskamp S, Cramer AOJ, Waldorp LJ, Schmittmann VD, Borsboom D. Qgraph: Network visualizations of relationships in psychometric data. J Stat Softw. 2012;48.

15. Friedman J, Hastie T, Tibshirani R. Sparse inverse covariance estimation with the graphical lasso. Biostatistics. 2007:1–10.
